# Pyruvate-conjugation of PEGylated liposomes effectively enhances their uptake in retinal photoreceptors

**DOI:** 10.1101/2022.11.18.517040

**Authors:** Gustav Christensen, Yiyi Chen, Dileep Urimi, Nicolaas Schipper, François Paquet-Durand

## Abstract

Despite several promising candidates there is a paucity of drug treatments available for patients suffering from retinal diseases. An important reason for this is the lack of suitable delivery systems that can achieve sufficiently high drug uptake in the retina and its photoreceptors. A promising and versatile method for drug delivery to specific cell types involves liposomes, surface-coated with substrates for transporter proteins highly expressed on the target cell.

We identified strong lactate transporter (monocarboxylate transporter, MCT) expression on photoreceptors as a potential target for drug delivery vehicles. To evaluate MCT suitability for drug targeting, we used PEG-coated liposomes and conjugated these with different monocarboxylates, including lactate, pyruvate, and cysteine. Monocarboxylate-conjugated dye-loaded liposomes were tested on both human-derived cell-lines and murine retinal explant cultures. We found that liposomes conjugated with pyruvate consistently displayed higher cell uptake than unconjugated liposomes or liposomes conjugated with lactate or cysteine. Pharmacological inhibition of MCT1 and MCT2 reduced internalization, suggesting an MCT-mediated uptake mechanism. Pyruvate-conjugated liposomes loaded with the drug candidates CN03 and CN04 reduced photoreceptor cell death in murine *rd1* and *rd10* retinal degeneration models.

Overall, this study proposes pyruvate-conjugated liposomes as a vehicle for drug delivery specifically to photoreceptors. Notably, in retinal degeneration models, free drug solutions could not achieve the same therapeutic effect. Our study thus highlights pyruvate-conjugated liposomes as a promising system for drug delivery to retinal photoreceptors, as well as other neuronal cell types displaying high expression of MCT-type proteins.

## 1. Introduction

Inherited retinal degeneration (IRD) relates to a group of diseases, including retinitis pigmentosa and Leber’s congenital amaurosis, characterized by the progressive loss of photoreceptors, which ultimately leads to blindness [1–3]. Typically, IRD-type diseases display a primary loss of rod photoreceptors, which are responsible for vision under dim light conditions. Accordingly, initial disease symptoms include night-blindness. Once rods are lost, the cone photoreceptors, which mediate color and high acuity vision under daylight conditions, also degenerate, ultimately leading to complete blindness. To date, IRD-type diseases remain essentially untreatable, creating a high need for new therapeutic developments.

In many types of IRD, high levels of cyclic guanosine monophosphate (cGMP) in rod photoreceptors, caused for instance by the impairment of phosphodiesterase 6 (PDE6), are found [3]. PDE6 regulates cGMP levels and dysregulation can cause cGMP to reach pathological concentrations and over-activate important cGMP-dependent proteins, eventually leading to cell death [4]. To mitigate this effect, drug candidates, which are inhibitory analogues to cGMP (**Figure 1**), have been shown to promote survival of photoreceptors in mouse models of IRD [5]. Rescuing rod photoreceptors can provide functional protection of cone photoreceptors [6].

**Figure 1:**
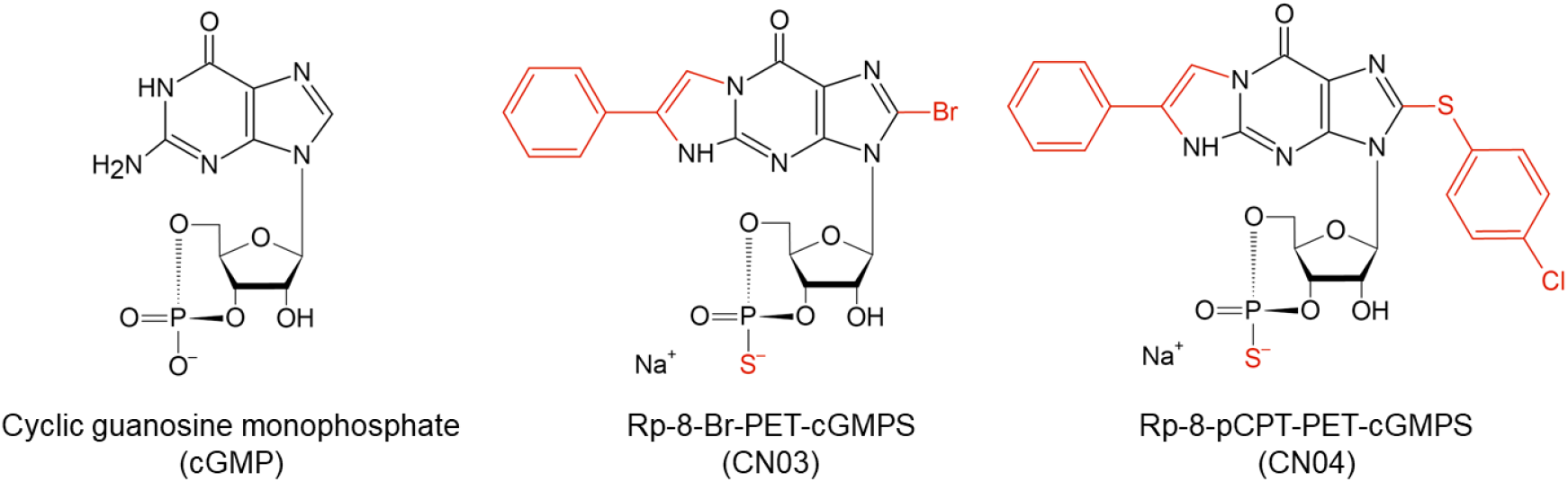
Chemical structures of the neuroprotective cGMP analogues CN03 and CN04. Modifications to the cGMP backbone are shown in red. Mw of CN03 = 562 g/mol. Mw of CN04 = 626 g/mol. For more information about these compounds please refer to [5].

To ensure a successful translation for clinical use, the cGMP analogues will need to be delivered to photoreceptors in high enough concentrations to provide a protective effect. This requires a suitable drug delivery system. Glutathione-conjugated liposomes were previously shown to enhance the therapeutic effect of CN03 in IRD mouse models after systemic administration [5]. These liposomes were designed for drug permeation across the blood-retinal barrier [7, 8]. However, intravitreal (IVT) administration may be preferable to achieve high drug concentrations at the target and to limit systemic exposure [9, 10]. The goal of this study was to design a liposomal system for IVT administration that was capable of directly targeting the photoreceptors to obtain a high therapeutic effect.

To achieve photoreceptor targeting, we decided to prepare liposomes conjugated with substrates for transporters expressed on photoreceptors. Other transporter-targeting liposomes and nanoparticles exist in the literature for delivery to different cell types [11–14]. Monocarboxylate transporters (MCTs) are membrane-bound transporters highly expressed in tissues which have a substantial energy demand and thus a high turnover of metabolites, including the retina [15]. MCTs are highly expressed on photoreceptors, possibly due to the demand for lactate shuttling between photoreceptors and Müller glial cells [16, 17]. Solid lipid nanoparticles conjugated with hydroxybutyric acid, a substrate for MCTs, have previously been prepared for brain targeting [18]. However, to our knowledge, no attempt has been made to target the photoreceptors in this manner. MCTs have a variety of substrates, including lactate, pyruvate, ketone bodies, and short-chain fatty acids [15, 19]. The most common substrates are lactate and pyruvate, and these were chosen as targeting ligand. Cysteine, which is structurally very similar to pyruvate, was also investigated to further pinpoint the exact changes in the chemical structures that are important for the uptake.

To test whether MCTs can mediate the uptake of monocarboxylate-liposomes, cell uptake in a human-derived cell-line (HEK293T) was investigated either with or without pretreatment with MCT inhibitors. Afterwards, organotypic retinal explant cultures derived from wild-type mice were used to determine the delivery of pyruvate-liposomes loaded with a hydrophilic dye to photoreceptors. Finally, the liposomal delivery of cGMP analogues to photoreceptors was determined by studying the effect of drug treatments in retinal cultures derived from IRD mouse models.

## 2. Methods

### 2.1 Animals

C3H *Pde6b^rd1/rd1^* (*rd1*), congenic C3H *Pde6b*^+/+^ wild-type (WT), and C57BL/6J *Pde6b^rd10/rd10^* (*rd10*) mice were housed under standard light conditions, had free access to food and water, and were used irrespective of gender. All procedures were performed in accordance with the association for research in vision and ophthalmology (ARVO) declaration for the use of animals in ophthalmic and vision research and the law on animal protection issued by the German Federal Government (Tierschutzgesetz) and were approved by the institutional animal welfare office of the University of Tübingen. All efforts were made to minimize the number of animals used and their suffering. Animals were not assigned to experimental groups prior to their sacrifice.

### 2.2 Materials

1-palmitoyl-2-oleoyl-sn-glycero-3-phosphocholine (POPC), distearoylphosphatidylcholine (DSPC), 1,2-distearoyl-sn-glycero-3-phosphoethanolamine-N-[methoxy(polyethylene glycol)-2000] (mPEG), 1,2-distearoyl-sn-glycero-3-phosphoethanolamine-N-[maleimide(polyethylene glycol)-2000] (maleimide-PEG), cholesterol, chloroform (99 % with 0.5-1 % ethanol), tris(2-carboxyethyl)phosphine (TCEP), thiolactate (95 %), L-cysteine, sodium mercaptopyruvate dihydrate, free calcein, hydrogen chloride, sodium hydroxide, disodium hydrogen phosphate dihydrate, sodium dihydrogen phosphate monohydrate, paraformaldehyde, Triton-X, goat serum, 4-(2-hydroxyethyl)-1-piperazineethanesulfonic acid (HEPES), sodium chloride, proteinase K, fetal bovine serum (FBS), bovine serum albumin (BSA), Corning™ Transwell polycarbonate membranes (0.4 μm, sterile), polycarbonate membranes (0.1 μm pore size), and filter supports, were obtained from Sigma-Aldrich (Darmstadt, Germany). Dulbecco’s modified eagle medium (DMEM), 1 % penicillin/streptomycin, R16 medium and dialysis cassettes (Slide-A-Lyzer, cellulose, 100K molecular weight cut-off) were obtained from Thermo Fisher Scientific (Waltham, MA, USA). The drug compounds CN03 and CN04 were provided by Biolog Life Science Institute (Bremen, Germany).

### 2.3 Preparation of monocarboxylate-liposomes

Liposomes with or without conjugation of mono-carboxylate-carrying molecules were prepared using the thin-film rehydration method. Here, a chloroform solution of the lipids POPC, cholesterol, and mPEG were mixed in a molar ratio of 63.3:31.7:5 to produce untargeted liposomes with the end of the PEG chain consisting of a methoxy group (Lp-OMe). Conversely, the lipids POPC, cholesterol, and PEG-maleimide were mixed in the same ratio to produce liposomes with a maleimide group at the end of the PEG chain for subsequent surface conjugation. For CN03 encapsulation, the lipid DSPC was used instead of POPC. All lipid solutions were dried with a rotation evaporator (KNF Neuberger, Trenton, NJ, USA) under reduced pressure (300 mbar) at 105 rpm and room temperature. After 1 h, the dried lipids were rehydrated in either of the rehydration buffers listed in **table 1**. For CN03 and CN04 encapsulation, the drugs were added in a 1:3 drug-to-lipid molar ratio. After lipid dissolution in the rehydration medium, 5 freeze-thaw cycles were performed in liquid nitrogen and 37 °C water bath. The liposome solutions were extruded at least 11 times through a PC membrane with 100 nm pores. For conjugation with monocarboxylate molecules, either thiolactate (for lactate-liposomes, Lp-Lac), sodium mercaptopyruvate (for pyruvate-coated liposomes, Lp-Pyr) or L-cysteine (for cysteine-coated liposomes, Lp-Cys) were prepared in 10 mM of the reducing agent tris(2-carboxyethyl)phosphine in 25 mM HEPES (pH 7.4) and added to PEG-maleimide-containing liposomes at twice the maleimide concentration followed by incubation at room temperature for 2 h. To remove non-encapsulated compounds and unbound monocarboxylates, the formulations were dialyzed against isotonic saline at 4 °C. For removal of drugs, a 2 h dialysis period was performed. For removal of calcein, a 6 h dialysis period was used with saline exchanged every second hour. The liposomes were sterile filtered and stored at 2-8 °C until further use. The drug concentrations were measured with Ultra High Performance Liquid Chromatography (UPLC) before and after dialysis. The hydrodynamic diameters and ζ-potentials were determined with dynamic light scattering (DLS) (see [20] for details).

**Table 1:** Solutions used for lipid rehydration in liposome preparation. All mediums adjusted to pH 7.4.

| Samples | Rehydration medium | Liposome concentration (mg/mL) |
| --- | --- | --- |
| Calcein-loaded liposomes | 2 mM calcein, PBS | 5 |
| CN03-loaded liposomes | 1.5 mM CN03, 25 mM HEPES, 125 mM NaCl | 5 |
| CN04-loaded liposomes | 281 μM CN04, 25 mM HEPES, 125 mM NaCl | 1 |
| Empty liposomes | PBS | 1 |
| Liposomes for ζ-potential measurements | 25 mM HEPES | 1 |

### 2.4 Uptake of monocarboxylate-liposomes in cell cultures

Cells of the human-derived cell line HEK293T were seeded in a 48-well plate at 50,000 cells/well in cell culture medium (DMEM + 20 % FBS + 1 % penicillin/streptomycin) for 24 h (37 °C, 5 % CO_2_). Calcein-loaded liposomes (Lp-OMe, Lp-Lac, Lp-Pyr, or Lp-Cys) were adjusted to 90 μM calcein and diluted 1:1 in the medium to give a final concentration of 45 μM. The same concentration of free calcein was added. To determine the role of MCTs in the uptake, some of the cells were pre-treated with either the MCT inhibitors AZD3965 (cat.no. HY-12750, MedChenExpress, New Jersey, USA) or AR-C155858 (cat.no. 4960, Tocris Bioscience, Bristol, UK) at a 2.5 μM and 1 μM concentration, respectively, for 24 h before calcein-loaded Lp-Pyr was added. For control, nothing was added to the cells. After a 2 h incubation period, the wells were washed three times with preheated (37°C) PBS. The fluorescent intensities (FI) were measured on a microplate reader (Spark 10M, Tecan, Männedorf, Switzerland) at Ex./Em. wavelengths of 485/530 nm to quantify the amount of intracellular calcein. The FI of cells without calcein were used as background signal. 90 μM calcein in PBS were measured to establish the total FI signal (100 %) for quantification. To prepare cells for imaging under fluorescent microscopy, sterile circular glass inserts (Ø 13 mm, Menzel-Gläser, Braunschweig, Germany) were added to wells of a 24-well plate before the cells were seeded for 24 h. After incubation with liposomes, the cells were fixed in 4 % paraformaldehyde for 15 min. The glass inserts were transferred to microscopy slides (Superfrost Plus^TM^, R. Langenbrinck, Emmendingen, Germany) and supplied with mounting medium containing DAPI (Vectashield, Vector laboratories, Burlingame, CA, USA). The cells were imaged on a fluorescent microscope (Axio Imager Z2, Zeiss, Oberkochen, Germany) with an ApoTome function with a CCD camera and a 20X objective. A green channel (Ex./Em. 493/517 nm) was used to measure the calcein signal, and a blue channel (Ex./Em. 353/465) was used to measure the DAPI signal.

### 2.5 Immunostaining of HEK293T cells and murine retinas

Sterile circular glass inserts were added to a 24-well plate before seeding with HEK293T cells (100,000 cells/well). After 24 h, the cells were fixed in 4 % paraformaldehyde and incubated with 0.3 % Triton-X in PBS for 5 min at room temperature followed by three times PBS washing. Afterwards, 5 % goat serum were added for 1 h. Primary antibodies against MCT1-4 (see **table 2**) were diluted in 5 % goat serum, added to the cells, and incubated overnight at 2-8 °C, followed by three times washing. The secondary antibody dissolved in 5 % goat serum 1:350 was added and incubated with the cells for 1 h under room temperature and then washed.

**Table 2:** List of antibodies used in this study. ICC = immunocytochemistry, IHC = immunohistochemistry.

| Antibody | Type | Provider, Cat.no. | Dilution |
| --- | --- | --- | --- |
| Anti-MCT 1 | Rabbit polyclonal | Alomone Labs, AMT-011 | 1:150 (ICC)<br>1:100 (IHC) |
| Anti-MCT 2 | Rabbit polyclonal | Alomone Labs, AMT-012 | 1:150 (ICC)<br>1:100 (IHC) |
| Anti-MCT 3 | Rabbit polyclonal | Abcam, ab60333 | 1:150 (ICC)<br>1:100 (IHC) |
| Anti-MCT 4 | Rabbit polyclonal | Abcam, ab180699 | 1:150 (ICC)<br>1:100 (IHC) |
| Anti-Glutamine Synthetase | Mouse monoclonal | Chemicon, MAB302 | 1:300 (IHC) |
| Anti-cone arrestin | Rabbit polyclonal | Sigma-Aldrich, ab15282 | 1:200 (IHC) |

For immunostaining of murine retinas, wild-type mice were used at post-natal day (P) 30. They were sacrificed by CO_2_ asphyxiation and cervical dislocation. The eyes were enucleated and the retinas isolated and fixed in 4 % paraformaldehyde, followed by cryoprotection in sucrose as described under [21]. The retinas were submerged in embedding medium (Tissue-Tek O.C.T. Compound, Sakura Finetek Europe, Alphen aan den Rijn, Netherlands), and frozen with liquid N2. 12 μm thick sections on microscope slides were produced using a cryostat (NX50, ThermoFisher, Waltham, MA, USA). The slides were dried and hydrated with PBS for 10 min. A blocking solution (10 % goat serum, 1 % BSA, and 0.3 % Triton-X in PBS) was added to the slides for 1 h. Primary antibodies against MCT 1-4, glutamine synthetase (GS), and cone arrestin were dissolved in the blocking solution and added to the slides, which were incubated overnight at 2-8 °C. Afterwards, they were rinsed with PBS three times, incubated with the secondary antibody for 1 h in the dark, and washed again with PBS. Mounting medium with DAPI was applied, and the slides were imaged using fluorescent microscopy. For detection of MCT1 and GS, the following channel was used: Ex./Em. 557/572. For detection of MCT2, the channel was: Ex./Em. 577/603. For detection of MCT3 and MCT4, the channel was Ex./Em. 493/517.

### 2.6 Uptake of monocarboxylate-liposomes in organotypic retinal explant cultures

Retinas were isolated and cultured from wild-type mice at post-natal day (P) 13 following a previously established protocol [21] and kept in culture until P15, after which a drop of 20 μL 5 mg/mL calcein-loaded liposomes (Lp-OMe, Lp-Pyr, or Lp-Cys) was added to top of the culture on the side corresponding to the vitreoretinal interface (*i.e*. the ganglion cell side). Alternatively, 1 μM AR-C155858 was added to the organ culture medium before addition of Lp-OMe or Lp-Pyr. After a 6 h incubation period (37 °C, 5 % CO_2_), the retinal explant cultures were fixed in 4 % paraformaldehyde, cryoprotected in sucrose and frozen with liquid N2, following the above mentioned protocol. 14 μm sections of retinal explant cultures from the center of the tissue were obtained using a NX50 cryostat (Thermofisher). The sections were hydrated with PBS for 10 min, supplied with mountain medium with DAPI, and imaged with fluorescent microscopy to record the DAPI signal (Ex./Em. 353/465 nm) and calcein signal (Ex./Em. 493/517 nm). Z-stacks were obtained by recording 11 images 1 μm apart. The stacks were projected using the Maximum Intensity Projection (MIP) function. From these images, the fluorescent intensity was measured, using the acquisition software (ZEN 2.6, Zeiss, Oberkochen, Germany), for each of the following layers: ganglion cell layer, inner plexiform layer, inner nuclear layer, outer plexiform layer, outer nuclear layer, photoreceptor inner and outer segments. Additionally, an immunostaining for cone photoreceptors (cone-arrestin) was performed following the immunostaining procedure for murine retina sections mentioned above, on sections from cultures incubated with calcein loaded Lp-Pyr.

### 2.7 Therapeutic effects of liposome-delivered CN03 and CN04 to photoreceptors in retinal explant cultures

Retinas derived from the *rd1* or *rd10* mouse models were cultured following the protocol under [21] and treated with the drugs CN03 or CN04. The treatment paradigms are shown in **Figure 3**. The same paradigm was followed on non-treated cultures. A treatment was done by applying a 20 μL solution containing 160 μM drug (either loaded in liposomes or in a free solution) to the top of the cultures. Assuming an even distribution in the culturing medium the final drug concentration in the medium would correspond to 3.14 μM. After fixation with 4 % paraformaldehyde, the histological work-up described above was followed to produce 14 μm thick sections. On the *rd1* derived culture sections, a terminal deoxynucleotidyl transferase dUTP nick end labeling (TUNEL) assay was performed, detailed under a published protocol [22]. Mounting medium with DAPI was applied, and the sections imaged with fluorescent microscopy. For TUNEL detection, Ex./Em. 548/561 nm filters were used. 11 z-stacks 1 μm apart were recorded, and from the projected images, the number of TUNEL-positive cells in the outer nuclear layer (ONL) was manually counted and expressed as a percentage with the formula TUNEL-positive cells (%) = Number of labelled cells in ONL / [(area of ONL)/(average area of one cell)]. For the *rd10* derived cultures, the average number of photoreceptor rows residing in the outer nuclear layer was counted from microscopy images, using the same imaging method.

**Figure 2:**
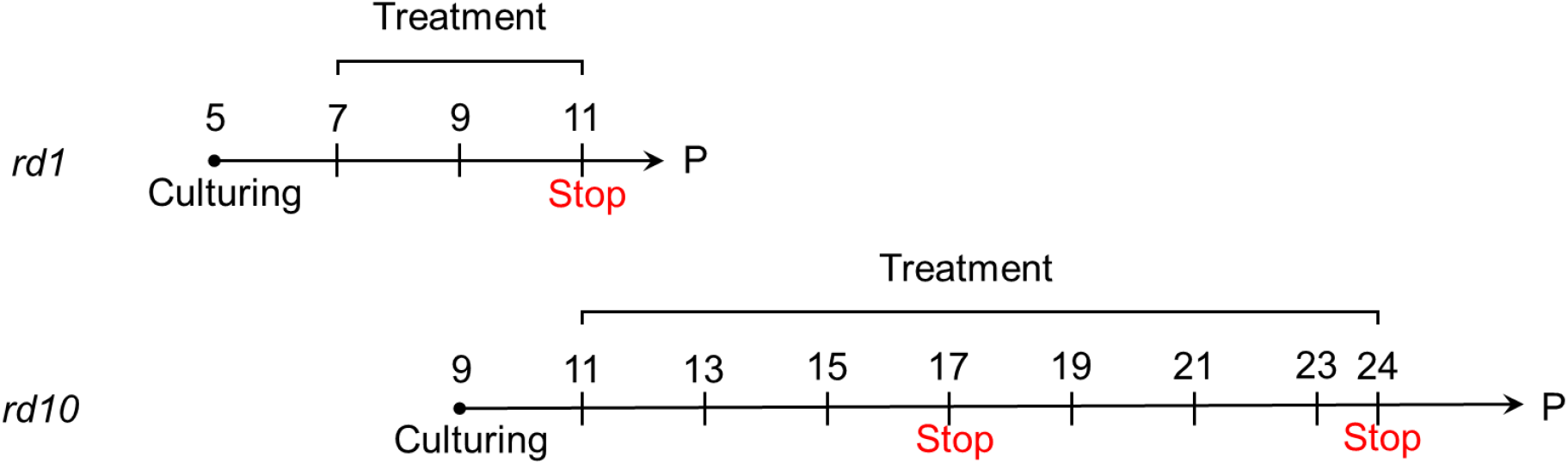
Treatment paradigms of organotypic retinal explant cultures derived from either the rapid degeneration *rd1* or the slow degeneration *rd10* mouse model. The cultures were treated every second day and stopped by chemical fixation at the indicated time-points. P = post-natal day.

## 3. Results

### 3.1 Expression of MCTs in the murine retina

We first investigated the expression of MCTs in the retina and specifically on photoreceptors to assess whether these transporters could be used for liposome targeting [18]. Immunostaining for MCT isoforms 1-4 was performed on retinal tissue sections (**Figure 3**). Different isoforms of MCTs were expressed in different areas of the retina. MCT1 and MCT2 were found to be expressed on photoreceptors. MCT1 was predominately localized to the inner segments of photoreceptors, while MCT2 was found on cell bodies in the outer nuclear layer. Due to their localization close to the outer border of the ONL, it is likely that MCT2 positive cells were cone photoreceptors. Most of the cells in the inner nuclear layer also expressed MCT2. MCT3 was not detected in the neuroretina. MCT4 was localized predominately at the vitreoretinal interface. A co-staining with the Müller glial cell marker glutamine synthetase revealed co-expression of MCT4, especially at the end-feet of Müller glial cells.

**Figure 3:**
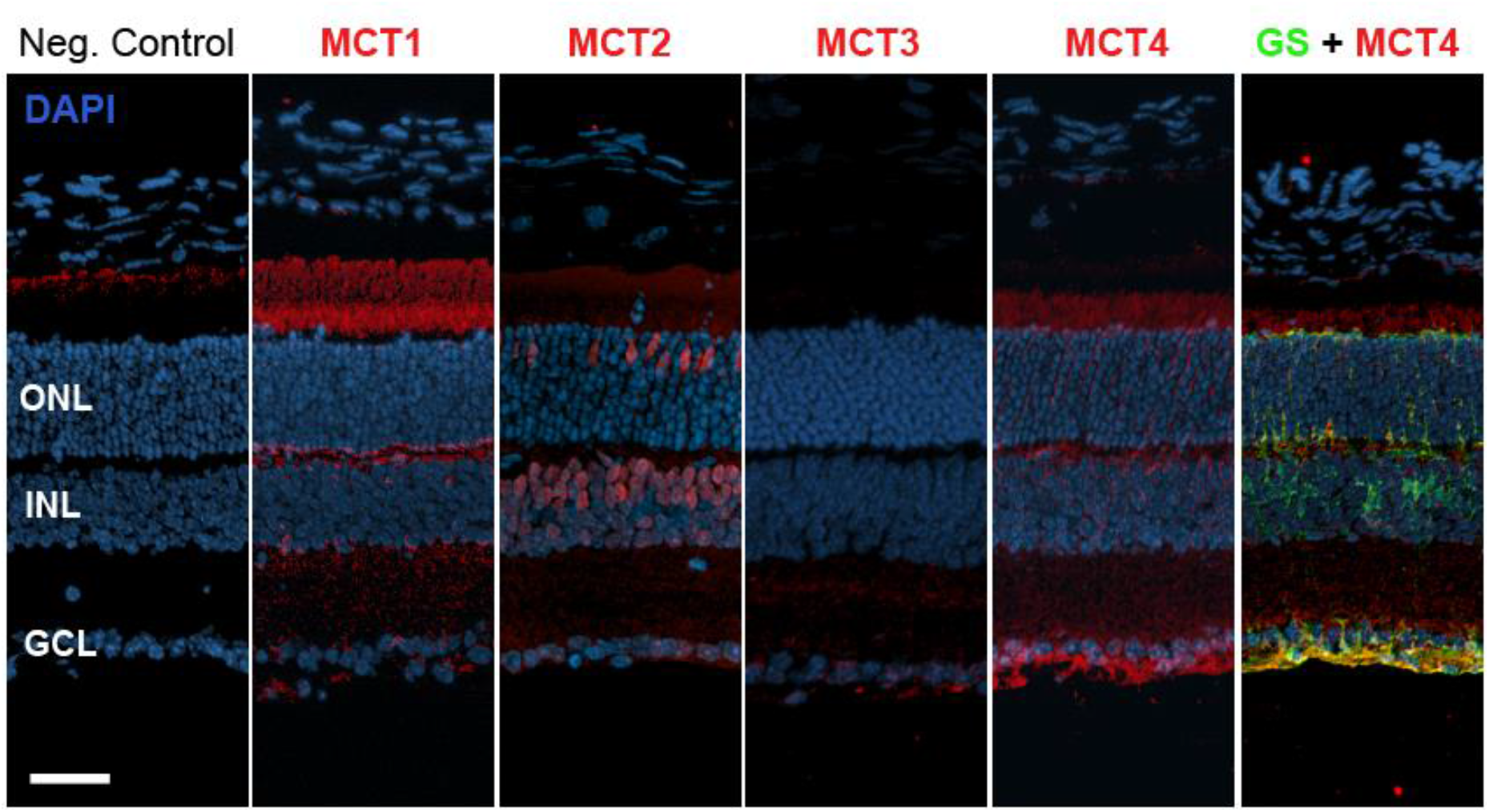
Immunostaining against monocarboxylate transporters (MCTs) in the retina. Expression of MCT isoforms 1-4 (red) in retinal sections obtained from wild-type mice at P30. Glutamine synthetase (GS) staining (green) was used as a marker for Müller glial cells and co-localized with MCT4 (yellow) notably in Müller cell end-feet. DAPI was used as nuclear counterstain (blue). Negative control is for the red channel only. GCL = ganglion cell layer, INL = inner nuclear layer, ONL = outer nuclear layer. Scale bar = 50 μm.

### 3.2 Characterization of monocarboxylate-liposomes

Since MCTs were expressed on retinal photoreceptors, we wanted to design a liposome system conjugated with substrates for MCTs. Lactate and pyruvate are common substrates for MCTs, so liposomes coupled with these molecules were prepared. Cysteine is structurally similar to both molecules, but is not considered a substrate of MCTs [19] and liposomes conjugated with cysteine may provide a passively transported control. All molecules were conjugated to the end of poly(ethylene glycol) (PEG) linked to the liposome surface. To confirm the successful formation of liposomes, dynamic light scattering (DLS) was performed to measure their hydrodynamic diameter and ζ-potential (**Figure 4**). The size of conjugated liposomes was the same as that of untargeted, control liposomes. The pyruvate and lactate liposomes showed a more negative surface potential than the control liposomes, indicating a successful conjugation. Since cysteine is neutral at pH 7, the cysteine-liposomes displayed the same ζ-potential as the control.

**Figure 4:**
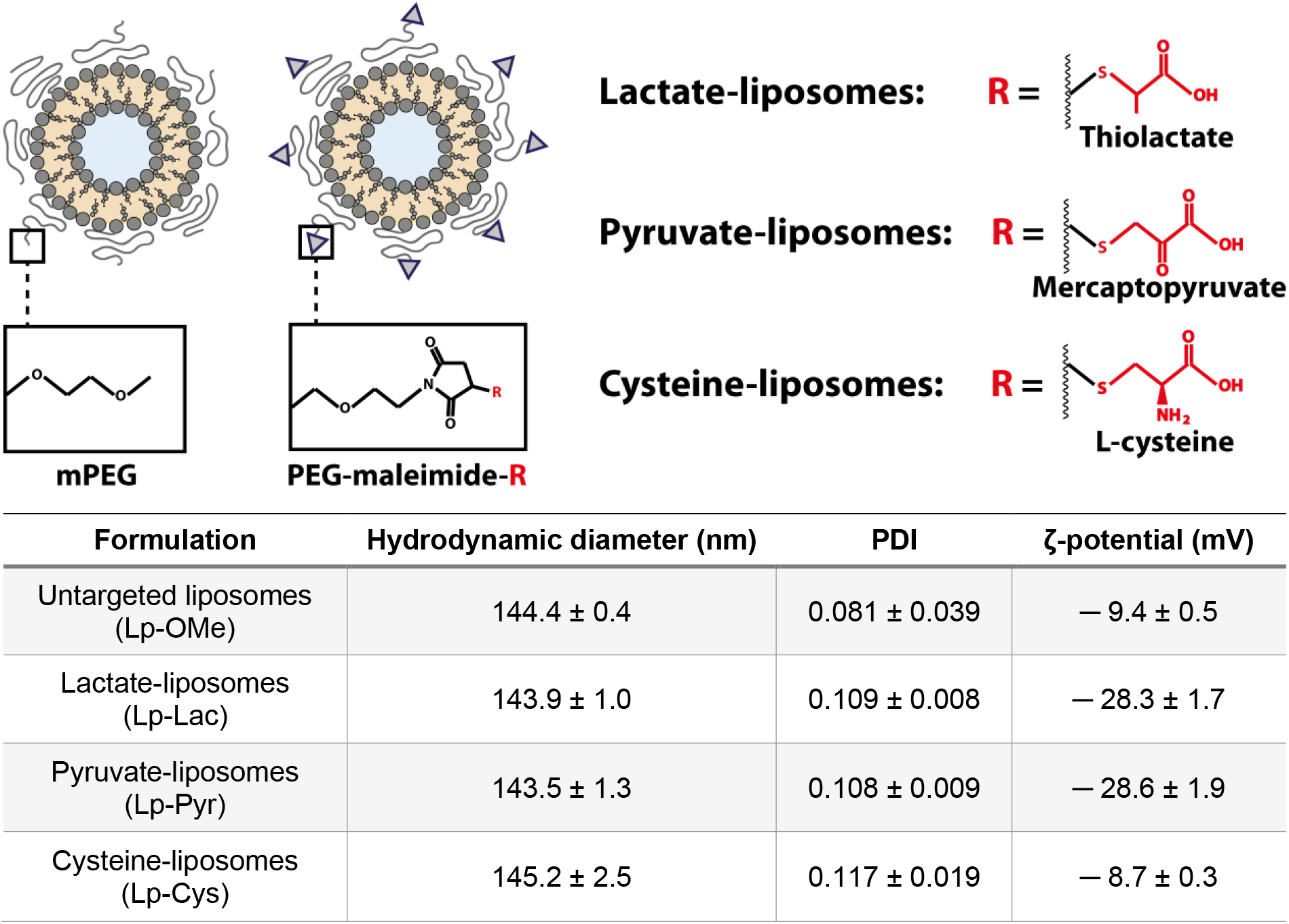
General structure and physical properties of liposomes prepared for this study. Liposomes were conjugated with different monocarboxylates at the end of surface-grafted poly-ethylene-glycol (PEG)-chains. Each monocarboxylate (R) was conjugated to PEG through a maleimide linker by a covalent attachment. Untargeted liposomes with methoxy-PEG (mPEG) were used as control. Hydrodynamic diameter, polydispersity index (PDI), and ζ-potential were quantified with dynamic light scattering. Data represent mean ± SD for n = 3.

### 3.3 Cellular uptake of monocarboxylate-coated liposomes

As a first proof-of-concept, the cellular uptake of monocarboxylate-coated liposomes was investigated using a human embryonic kidney cell line (HEK293T). An immunostaining demonstrated the expression of MCT isoforms 1-4 in these cells (**Figure 5A**). All liposome formulations (either coupled with monocarboxylate or untargeted) were loaded with calcein, which is a cell impermeable green fluorescent dye, and incubated with the cells for 2 h. Fluorescent images revealed high uptake of pyruvate- and cysteine-coated liposomes compared to the other conditions (**Figure 5B**). Calcein uptake in the cells was significantly increased with pyruvate- and cysteine-liposomes (**Figure 5C**), with the former displaying higher uptake than the latter. Cysteine-coated nanoparticles have previously been shown to be taken up HEK293 cells [23].

**Figure 5:**
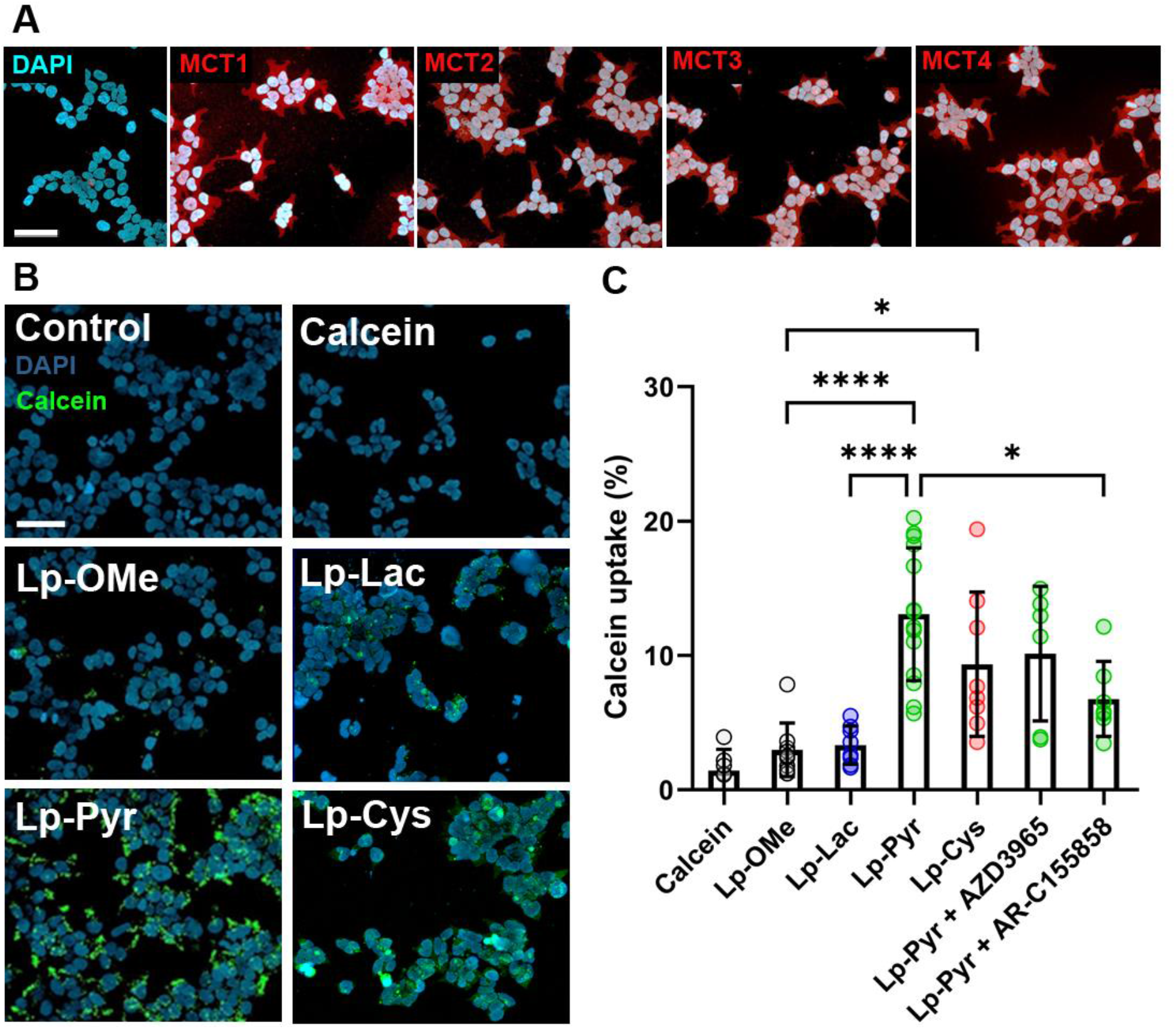
Cellular uptake of monocarboxylate-coupled liposomes. **A** Immunostaining against MCT1-4 isoforms in HEK293T cells. Red: immunostaining, blue: DAPI. **B** Fluorescent microscopy images of cell cultures after incubation with monocarboxylate-liposomes. **C** Amount of internalized calcein relative to total added calcein. Results presented as mean ± SD, * = p ≤ 0.05, **** = p ≤ 0.0001. Scale bars = 50 μm.

Interestingly, when specific inhibitors against MCTs (AZD3965 and AR-C155858) were added, the pyruvate-liposomes delivered less calcein to the cells, suggesting that the uptake was at least partially mediated by MCTs. When the inhibitor AR-C155858 was used, this difference was significant. Since AZD3965 is selective towards MCT1 (Ki = 1.6 nM [24]), while AR-C155858 inhibits both MCT1 and MCT2 (Ki = 2.3 and <10 nM, respectively [25]), this suggests that the pyruvate-liposomes might be more prone to MCT2-dependent uptake. Even when the AR-C155858 inhibitor was added, there was still a tendency towards higher calcein uptake by Lp-Pyr compared to Lp-OMe. This suggests that in HEK293 cells also MCT3 or MCT4 could mediate uptake. Both transporters have been found to actively take up lactate *in vitro* [26, 27].

To demonstrate that the enhanced cell uptake of pyruvate-conjugated nanoparticles was not restricted to liposomes, we tested the uptake of micelles produced from PEG-conjugated lipids (**Figure S1**). The uptake of micelles with or without pyruvate-conjugation loaded with a hydrophobic dye were tested in HEK293T cells, and similar results compared with the liposome uptake were obtained.

### 3.4 Retinal uptake of pyruvate-coated liposomes

To determine the potential of pyruvate-liposomes to deliver drug to photoreceptors, their uptake in organotypic retinal explant cultures derived from mice were analyzed. Since the target delivery compounds are hydrophilic molecules, a similarly hydrophilic dye, calcein, was used to predict where in the tissue the drugs would accumulate. Calcein was loaded into either the MCT-targeting pyruvate-liposomes (Lp-Pyr), cysteine-liposomes (Lp-Cys), or untargeted control liposomes (Lp-OMe). These were added to retinal explant cultures at post-natal day 15 to the vitreous-facing side of the isolated retinas to simulate the IVT route. After an incubation period of 6 h, the cultures were fixed, frozen, and sectioned. The amount of calcein dye in the sections was analyzed from fluorescent microscopy images (**Figure 6A**) and the signal measured from the distinct retinal layers (**Figure 6B**). Lp-Pyr achieved more calcein uptake compared to Lp-OMe in the inner plexiform layer (IPL) and in the outer retina, from the outer plexiform layer (OPL) to the photoreceptor segments. More calcein signal for Lp-Pyr than for Lp-Cys could be detected in the ONL. This comparison is especially relevant since the two formulations share very similar structures and sizes (*cf*. **Figure 4**).

**Figure 6:**
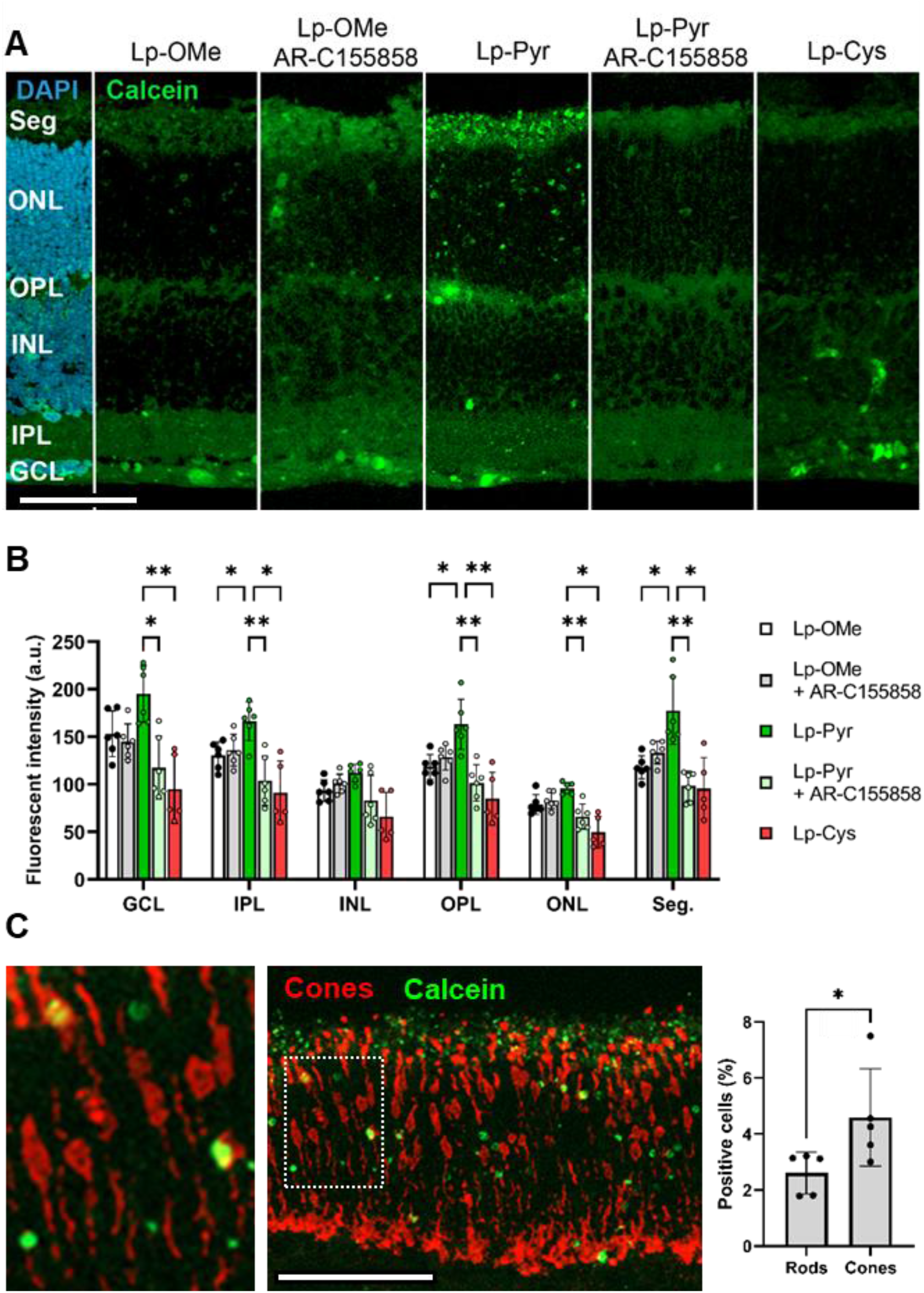
Distribution of liposome-delivered calcein in organotypic retinal explant cultures. Monocarboxlate-liposomes (Lp-Pyr and Lp-Cys) containing the hydrophilic dye calcein were added to retinal cultures to the side closest to the ganglion cell layer (GCL) at post-natal day 15 for 6 h and compared to untargeted liposomes (Lp-OMe) and in combination with the MCT1-2 inhibitor AR-C155858. **A** Representative images demonstrating calcein distribution in the retina. IPL = inner plexiform layer, INL = inner nuclear layer, OPL = outer plexiform layer, ONL = outer nuclear layer, Seg. = photoreceptor inner and outer segments. Green = calcein signal. Blue = DAPI nuclear counterstain. **B** Signal from the green channel for each retinal layer. Results represent mean ± SD for n = 5-6, * = p ≤ 0.05, ** = p ≤ 0.01. Statistical analysis: Two-way ANOVA with Tukey’s multiple comparison test. **C** Immunostaining of cone-arrestin as a cone marker in sections from retinal cultures to which calcein-loaded Lp-Pyr was added. The number of rods and cones that contained detectable amounts of calcein in the ONL was analyzed. Results represent mean ± SD for n = 5, * = p ≤ 0.05. Statistical analysis: two-tailed unpaired t-test. Scale bars = 50 μm.

We also investigated the intracellular distribution in photoreceptor from close-up images (see **Figure S2**), and after a qualitative analysis, we found that calcein accumulated in the outer segment and cell body of photoreceptors. This is after a 6 h incubation period, and it is likely that after longer time-periods, calcein would distribute evenly in the cells due to its small molecular weight. However, the results are relevant for the delivery of high molecular weight compounds like peptides or oligonucleotides with an intracellular target.

To test if the higher uptake was mediated by MCTs, the retinal cultures were treated with the MCT1-2 inhibitor AR-C155858 during the incubation period with Lp-Pyr. When the inhibitor was added, an overall decrease in the calcein signal was observed, suggesting that the uptake was mediated by MCTs. Because AR-C155858 is not considered to be an inhibitor for MCT4 [25], our results indicate that MCT1-2 are especially important for Lp-Pyr uptake.

Since MCTs regulate the flow of metabolites in and out of the cells, the inhibition of these key transporters may restrict or slow down the cells’ capabilities of taking up liposomes due to low energy conditions or toxic side-effects. To control for this, Lp-OMe was used in combination with the MCT inhibitor. We found that, unlike the Lp-Pyr uptake, the addition of AR-C155858 did not lead to any reduction of Lp-OMe uptake, suggesting a direct link between the transporters and Lp-Pyr uptake.

From the MCT immunostaining results (**Figure 5**), it appeared that cones predominately express MCT2. Since MCT2 is thought to be more specific for pyruvate transport [15], we tested whether Lp-Pyr was delivered primarily to cones or rods in the outer retina. An immunostaining against the cone specific marker cone-arrestin was performed on sections from cultures to which Lp-Pyr was applied, and the relative amount of calcein-containing rods and cones were determined (**Figure 6C**). We found proportionally more cone uptake than rod uptake to a significant extend, a finding that could be relevant in the context of cone-specific diseases like achromatopsia or age-dependent macular degeneration (AMD).

### 3.5 Treatment effect of drug-loaded pyruvate-liposomes

Since the pyruvate-liposomes showed superior photoreceptor uptake, we next tested whether these liposomes could enhance the delivery of specific drugs to photoreceptors. Retinal explant cultures derived from the photoreceptor-degeneration mouse model *rd1* were used and treated from P7 to P11, *i.e*., a time-point just before the peak of degeneration. Both free CN03 and CN04, as well as encapsulated compounds were used to assess the effect of the liposome system. The TUNEL assay was employed to detect dying photoreceptors in tissue cross-sections as a read-out of the effect of different treatments (**Figure 7**).

**Figure 7:**
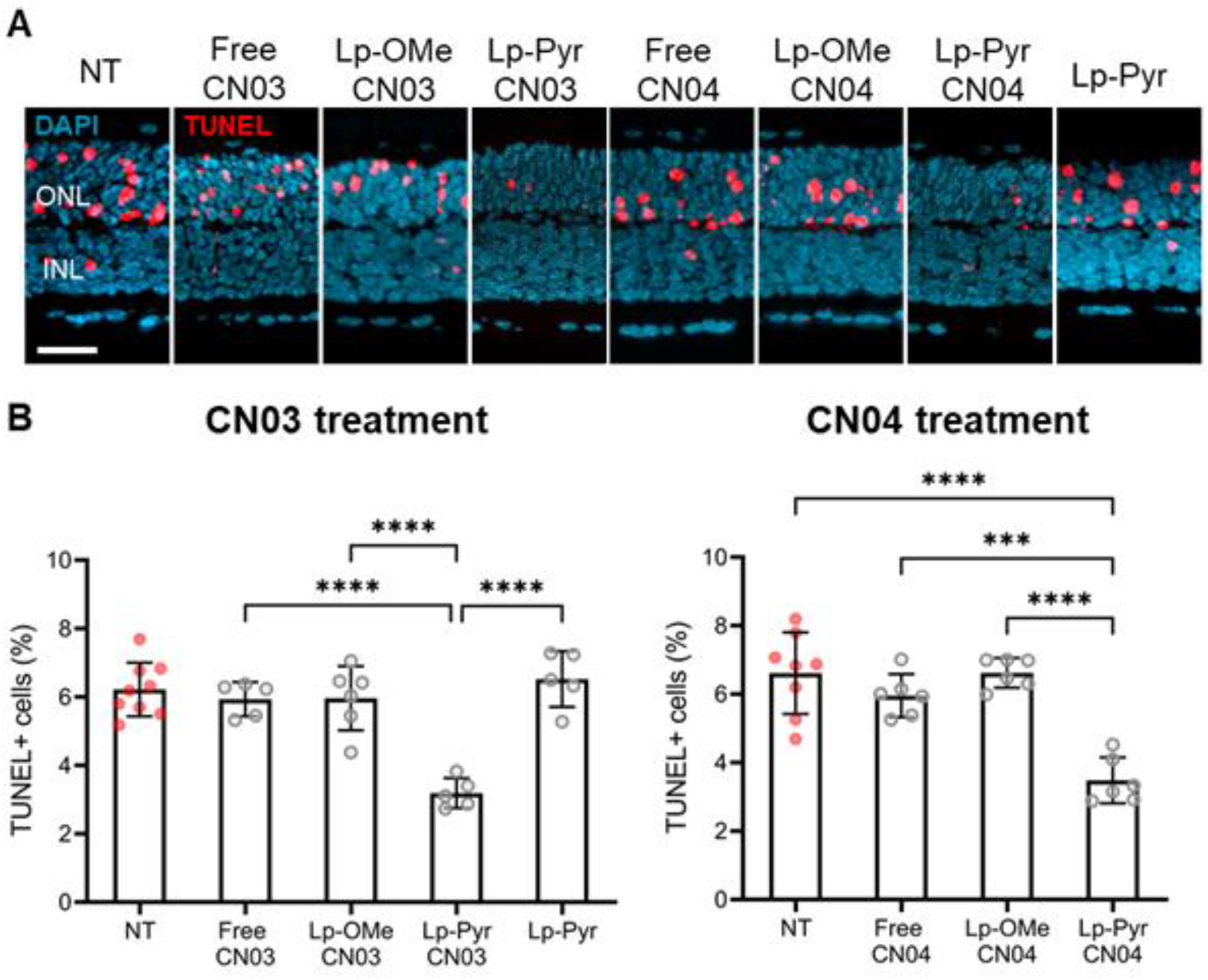
Treatment of organotypic retinal explant cultures derived from the *rd1* mouse model. Retinas were cultured at P5 and treated with either liposome-encapsulated or non-encapsulated (free) drugs (CN03 or CN04) from P7-P11. Assuming an equal distribution across the entire culturing medium, the final drug concentration was 3.14 μM for all treatments. **A** The amount of dying photoreceptors in tissue sections from cultures under various treatment conditions was assessed using the TUNEL assay (red). DAPI (blue) was used to label nuclei in the outer nuclear layer (ONL). **B** Percentages of dying (TUNEL+) cells in the ONL. Under non-treated (NT) conditions, the number of TUNEL+ cells was high in the *rd1* model. Administration of free compounds CN03 or CN04 (3.14 μM total concentration) had no significant effect on cell death. The same was observed for untargeted, control liposomes (Lp-OMe) and targeted but empty pyruvate-conjugated liposomes (Lp-Pyr). Importantly, Lp-Pyr, when loaded with CN03 or CN04, produced significant photoreceptor protection. Data are presented as mean ± SD for n = 5-9 animals, *** = p ≤ 0.001, **** = p ≤ 0.0001. Statistical analysis: One-way ANOVA with Tukey’s multiple comparison. Scale bar = 50 μm.

When free CN03 or CN04 was applied to the *rd1*-derived cultures, no significant reduction in dying photoreceptors was observed, although there was a tendency of reduced cell death. While CN03 and CN04 showed protection in this model at concentrations of 10 – 50 μM [5], the amount used here (160 μM in 20 μL diluted in 1 ml medium = 3.14 μM) was too low to achieve a reduction of cell death when using the free drug solution alone. The Lp-OMe system proved insufficient to improve the transport of CN03 or CN04 to photoreceptors. However, the Lp-Pyr system loaded with either CN03 or CN04 could significantly protect the photoreceptors. About 50 % less cell death was observed compared to the non-treated control (NT). This protection was the result of an enhanced drug transport and not the pyruvate-liposome, since empty pyruvate-liposomes did not provide any photoreceptor protection. The lipid concentration used here was 2 mg/mL as this was the approximate concentration in the Lp-Pyr/CN03 samples. For Lp-Pyr, the encapsulation efficiency was 24.7 ± 6.5 % for CN03 and 80.0 ± 5.9 % for CN04. Due to the low encapsulation efficiency of CN03, these samples had the highest lipid concentration.

The rate of cell death in the *rd1* model is very fast with almost complete rod loss at P18 [2], which is not representative for most IRD patients. A somewhat better representation of the human disease situation is afforded by the slower degeneration *rd10* mouse model, in which the peak of photoreceptor cell death occurs around P20. Hence, we tested whether pyruvate-liposomes could achieve similar benefits in this model. Here, we tested CN03 encapsulated into liposomes containing DSPC, a formulation that causes slower drug release [28] and is expected to be better suited for *in vivo* applications. In the *rd10* model, we assessed longer-term photoreceptor survival by quantifying the number of photoreceptor rows remaining in the tissue at P17 and P24 (*i.e*., before and after the peak of degeneration) (**Figure 8**). *rd10* photoreceptor protection was observed when CN03 was encapsulated in pyruvate-liposomes, while the free drug at the same concentration (3.14 μM) did not achieve a rescue effect.

**Figure 8:**
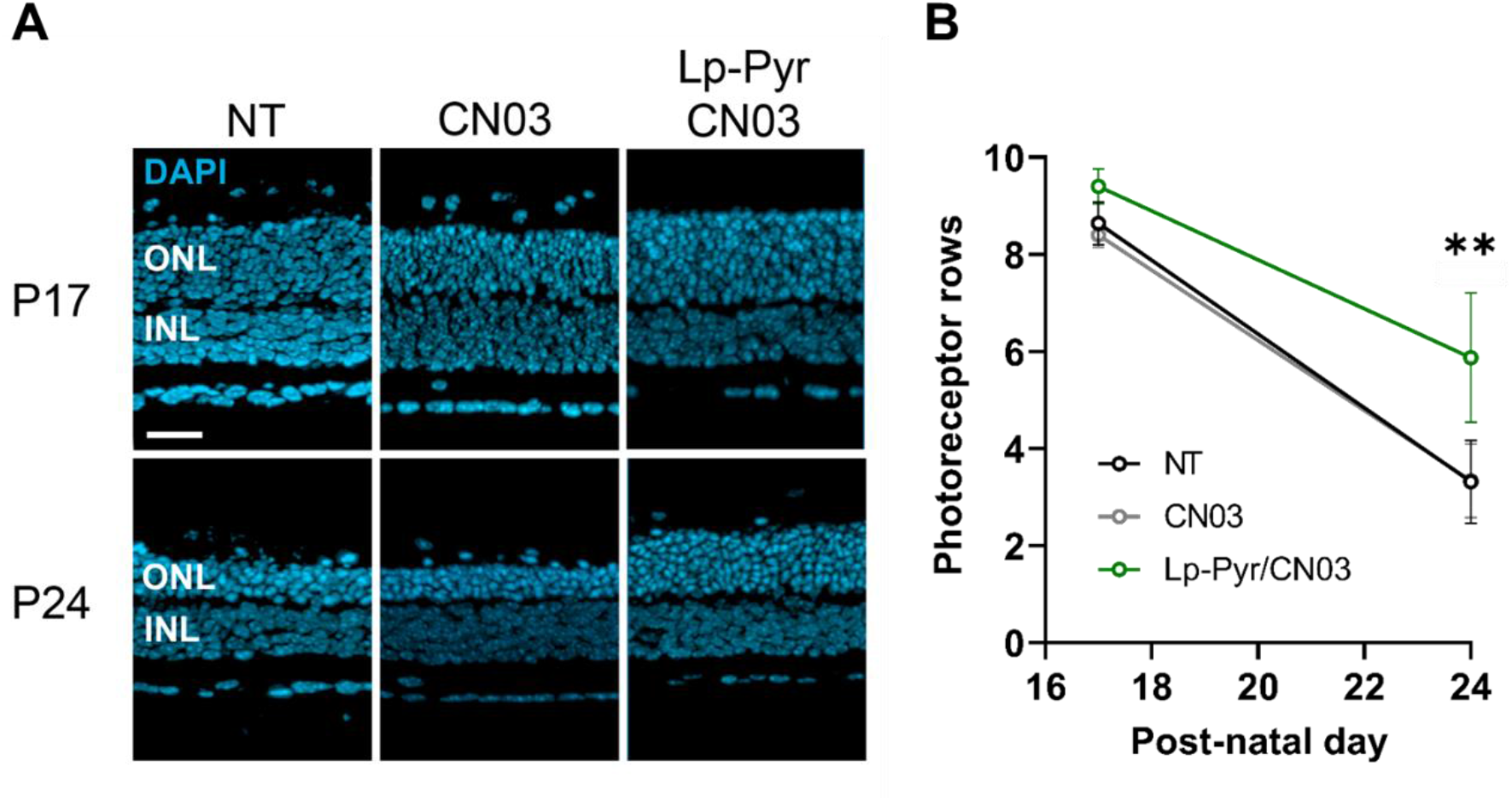
Treatment of organotypic retinal explant cultures derived from the *rd10* mice model. Retinas were cultured at P9 and treated from P11 until either P17 or P24 with free CN03 or with CN03 encapsulated in pyruvate-liposomes (Lp-Pyr/CN03). **A** Representative images of retinal explant culture sections at P17 and P24. DAPI (blue) as nuclear stain. ONL = outer nuclear layer, INL = inner nuclear layer. Scale bar = 50 μm. **B** Number of remaining photoreceptor rows in the tissue sections. Results represent mean ± SD. At P17: n = 4-6. At P24: n = 4-5. ** = p ≤ 0.01 between the Lp-Pyr/CN03 and CN03 or NT group. Statistical analysis: Two-way ANOVA with Tukey’s multiple comparison test.

## 4. Discussion

### 4.1 Pyruvate-liposome uptake in the retina and photoreceptors

Efficient retinal drug delivery is a critical concern, notably in the context of IRD. In this study, we have developed a novel approach to deliver cargo to photoreceptors using a targeted liposomal drug delivery system. We show that pyruvate-conjugated liposomes loaded with neuroprotective drugs can achieve efficient photoreceptor protection. While tested here for cGMP-analogue-based drugs, our approach could also benefit the delivery of biologics like peptides or genetic materials.

We found that when pyruvate was conjugated to liposome-grafted PEG-chains, more MCT-dependent cell uptake was observed in HEK293T cells and photoreceptor-uptake in retinal explant cultures. Furthermore, this photoreceptor-directed uptake improved the therapeutic effect of photoreceptor rescuing drugs *in vitro*. We attribute this effect to MCT1-2 transporters expressed on photoreceptors. Surprisingly, lactate-liposomes did not achieve the same benefit, possibly due to structural differences between lactate and thiolactate that was used in the preparation of lactate-liposomes (**Figure 4**). Thiolactate lacks the OH group present in native lactate recognized by MCTs. Thus, minor differences between the targeting ligand and native substrate may affect transport recognition. This is further exemplified by the fact that our cysteine-liposomes achieved poor uptake in the retina despite being structurally similar to pyruvate. In pyruvate-liposomes, mercaptopyruvate was used, which has a thiol group at the C3-position. The potent anti-cancer drug 3-bromopyruvate has been shown to target MCT1 expressed on cancer cells [29], illustrating how pyruvate modified at the C3-position can be recognized by MCTs.

For uptake in the retina, it is likely that the liposomes are taken up by Müller glial cells due to the phagocytotic nature of these cells [30]. Trans-retinal permeation of nanoparticles has been documented in the *in vivo* mouse retina [31, 32], which might occur by Müller cell transcytosis and subsequent release in the interphotoreceptor matrix [33]. Moreover, transporter-targeting can lead to improved transcytosis across tissue barriers such as the blood-brain barrier [8, 34] and the intestinal epithelium [12]. This suggests that pyruvate-liposomes can potentially penetrate the retina better than conventional liposomes, possibly via MCT4-mediated uptake into the Müller cells (Figure 9). In a future study, the Müller cell-transcytosis hypothesis, may perhaps be tested by using MCT4-specific inhibitors. Although photoreceptors are not as prone to endocytosis as other cells in the retina, like the Müller cells and RPE [35], one study found endocytic activity in the inner segments of photoreceptors [36]. The mechanism of liposome uptake by photoreceptors is beyond the scope of this study, however, several endocytic pathways for transporter-targeting nanoparticles have been reported in the literature [14]. Some studies found that the uptake was associated with the caveolae-mediated pathway, which avoids lysosomal digestion [13]. This is preferred for gene and peptide delivery. Another study found that both clathrin- and caveolae-mediated pathways were involved in the cellular uptake of vitamin B12-conjugated nanoparticles [11]. Not much information is available about the fate of the transporters following liposome uptake. One study found that the transporters were recycled on the cell surface after the initial liposome uptake [37], meaning that liposomes did not result in cell-triggered breakdown of the transporters.

**Figure 9:**
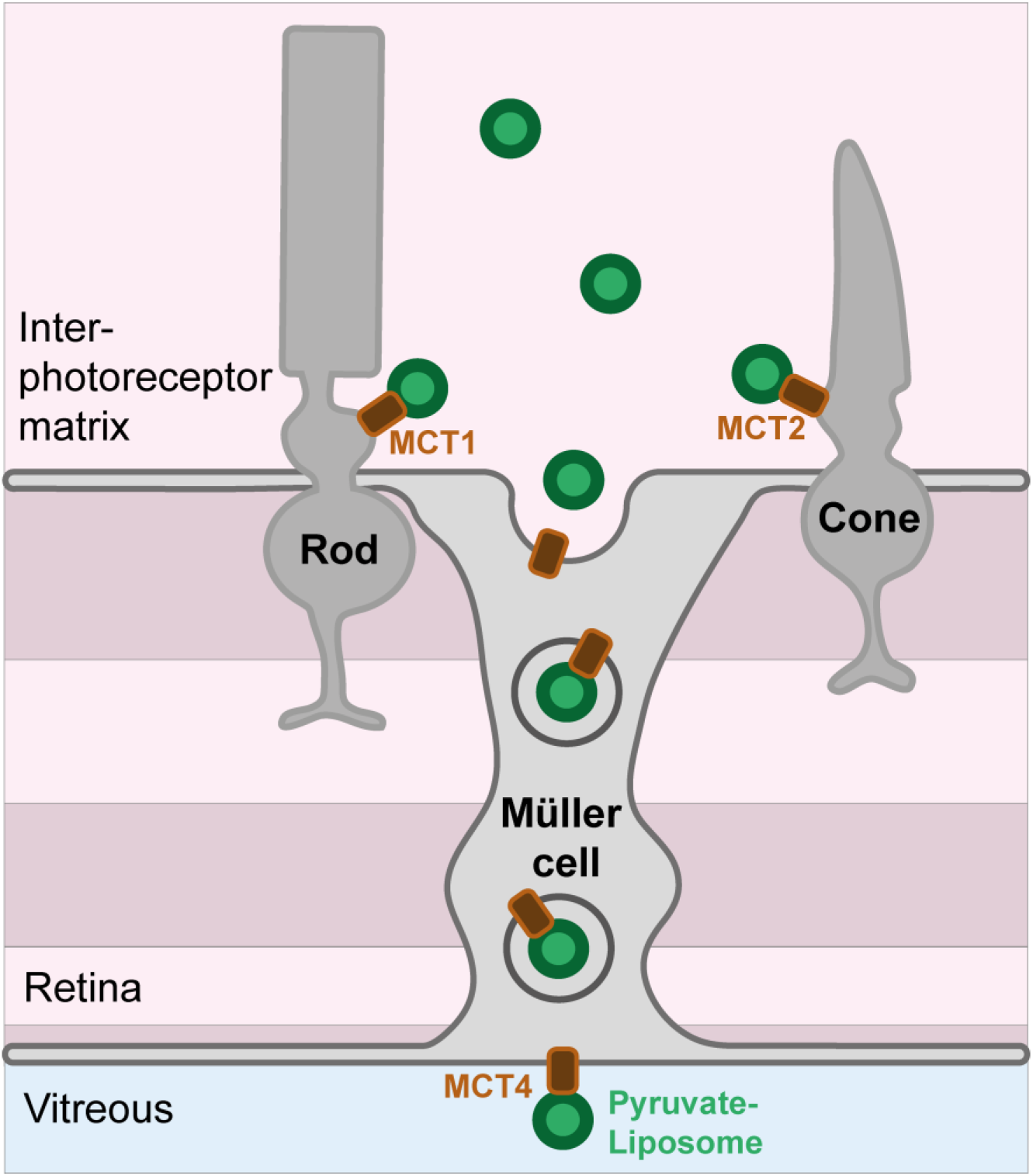
Potential mechanism for the transport of pyruvate-conjugated liposomes to photoreceptors. Müller cells via MCT4 may mediate the initital transport and release of liposomes into the inter-photoreceptor matrix. Here, a secondary uptake in rod or cone photoreceptors may be faclitated by either MCT1 or MCT2, respectively. MCT = monocarboxylate transporter.

### 4.2 Pyruvate-liposomes for clinical use

For our study, we have used murine explant retinal cultures, which have several advantages over *in vivo* models as they are cheaper, faster, more reliable, and, most importantly, allow studying direct, retina-specific effects. Yet, the rodent retina display differences compared with the human retina, especially regarding the vitreoretinal barriers. In bovine cultures, it was found that PEGylated liposomes less than 100 nm in diameter can reach the photoreceptors [38]. Since our liposomes are slightly larger than 100 nm, they may need to be shrunk down for permeation of the vitreoretinal barrier and future clinical use. Additional optimization of the pyruvate-liposomes may include using a shorter PEG chain [39]. In the explant culture system, we do not preserve the vitreous after the culturing, so the IVT mobility of the pyruvate-liposomes was not tested. Previously we have documented the bio-distribution of similar liposomes and found that the retina could be reached after an IVT injection into *ex vivo* porcine eyes [40]. This is in line with similar studies [38, 41].

In contrast to a free drug solution or untargeted drug delivery system, photoreceptor-targeting liposomes have the benefit of lowering the dosage required to achieve an effect because of higher cell uptake. This was demonstrated in this study for the drug candidates CN03 and CN04. In addition, higher dosage may be supplied since targeting would diminish unwanted side-effects in other cells and tissues. Both effects together could substantially widen the therapeutic window. This would make it easier to find a suitable dosing regimen and would allow for prolonged drug exposure in the target cell, minimizing administration frequency [42]. This in turn is especially relevant for IVT injections that carry a range of potential complications [40, 43–45].

How useful our photoreceptor-targeted liposomes are for clinical use remains to be studied and would, for example, depend on the drug release rate. Rapid drug release may limit the shelf-life of the formulation and can potentially be unsafe as injection of too much free drugs might cause toxic side-effects [46]. A high drug release rate also limits the use of the liposomes to a level that it is similar or equal to the effect of the free drug. For conventional liposomes, however, a very slow drug release rate can have inferior outcomes, because the drug is trapped in the liposome. Where the liposome cannot enter the target, the concentration of available drug in the target tissue could be too low to achieve an effect [47]. For targeted liposomes, conversely, it has been found that lower release rates are beneficial to the therapeutic effect of the drugs, [48]. The reason is likely due to the drug internalization in target cells, which was higher for slow drug-releasing liposomes. This suggests that there is in principle no limit in how slow the release rate can be for targeted liposomes. The currently studied liposomes may give rise to fairly rapid release of the drug. We have observed complete release of CN04 from similar liposomes in an *in vitro* set-up within 48 h (data not shown). This is comparable to the release rate of other hydrophilic compounds from liposomes [49]. The release rate for CN03 and CN04 may be lowered by precipitation within the liposome cavity with a specific salt using a remote loading technique similar to what has been done for liposomal doxorubicin formulations [50]. CN03 has previously been remote loaded in similar liposomes using calcium acetate salts [5]. Another strategy to lower the release rate would be to include a hydrophobic linker to the drugs. Hydrophobic drugs have a much slower release rate than hydrophilic drugs from liposomes [51]. Hence, the liposomal delivery system and the encapsulated drug together need to be tailored for the specific purpose and release profile. Such optimization of liposomes will also entail testing in live animal models to obtain the required pharmacokinetic datasets before clinical use can be envisaged.

Overall, our study indicates that pyruvate-liposomes are useful as a drug delivery system for the active targeting of photoreceptors. Combined with a suitable drug, this delivery approach may be very promising for the development of new treatments for chronic retinal diseases, notably for IRD and possibly also for cone degeneration diseases such as achromatopsia or AMD.

## 6. Acknowledgements

We would like to thank Dr. Pieter Gaillard for his useful insight into the topic and helpful suggestions regarding experimental design and manuscript editing and Pietro De Angeli for assistance with the HEK293T cells.

## 8. Author contributions

Conceptualization: G.C., F.P.-D.; Methodology: G.C., Y.C., F.P.-D.; Formal analysis: G.C., Y.C., D.U.; Investigation: G.C., Y.C., D.U.; Resources: F.P.-D., N.S.; Writing - Original Draft: G.C.; Writing - Review & Editing: G.C., Y.C., D.U., N.S., F.P.-D.; Visualization: G.C., Y.C.; Supervision: F.P.-D., N.S.; Funding acquisition: F.P.-D., N.S.

