## Supplementary material for "Pyruvate-conjugation of PEGylated liposomes effectively enhances their uptake in retinal photoreceptors": SI _ Pyruvate-conjugation of PEGylated liposomes effectively enhances their uptake in retinal photoreceptors.pdf

### Uptake of pyruvate-micelles in HEK293T cells

Micelles were prepared by dissolving 0.9 mg of either 1,2-distearoyl-sn-glycero-3-phosphoethanolamine-N-[maleimide(polyethylene glycol)-2000] (DSPE-PEG-maleimide) or 1,2-distearoyl-sn-glycero-3-phosphoethanolamine-N-[methoxy(polyethylene glycol)-2000] (DSPE-mPEG) together with 10 µg 3,3'-dioctadecyloxacarbocyanine perchlorate (DiO) in 99% chloroform with 0.5-1 % ethanol. The solvent was dried with a rotary evaporator (KNF Neuberger, Trenton, NJ, USA) for 1 h under 300 mbar at 120 rpm under room temperature. The dried films were hydrated in PBS to produce micelle solutions with a final concentration of 0.5 mM lipids and 20 µM DiO. To micelles consisting of DSPE-PEG-maleimide, sodium mercaptopyruvate dissolved in 10 mM tris(2-carboxyethyl)phosphine (pH 7.4) were added at a 1:2 maleimide-to-mercaptopyruvate mole ratio and left overnight to produce pyruvate-micelles. Micelles consisting of DSPE-mPEG were used as control. The DiO signal was measured on a microplate reader (Spark 10M, Tecan, Männedorf, Switzerland) using Ex./Em. wavelengths of 470/515 nm, and all solutions were diluted to have the same fluorescent intensities. HEK293T cells were seeded on a 96-well plate at a concentration of 25,000 cells/well overnight. Micelles containing DiO were added to the cells at a 1:1 dilution in the cell culture medium and incubated for 2 h (37 °C, 5 % $CO_2$). The cells were washed at least three times with preheated PBS. The fluorescent intensity of the DiO signal was measured, and signal from cells without micelles were used to determine the background signal. Signal from micelle stock solutions was used to quantify the DiO uptake.

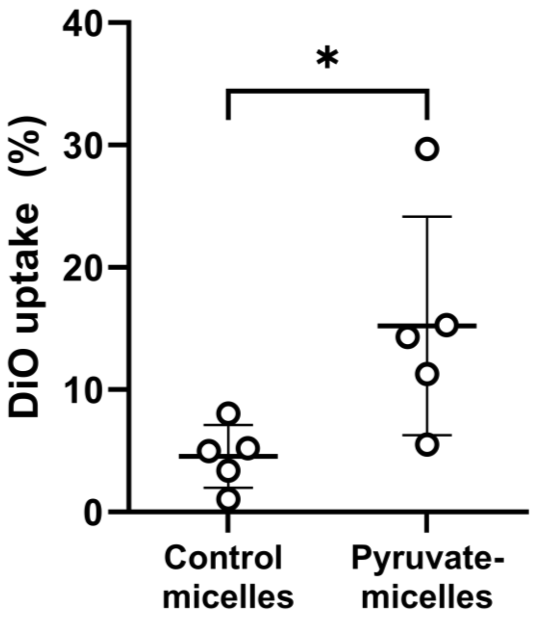


**Figure S1: Uptake of pyruvate-conjugated micelles in a human-derived cell line.** HEK239T cells incubated with lipid-PEG micelles with either methoxy-PEG (control micelles) or pyruvate-conjugated PEG (pyruvate-micelles) encapsulating the fluorophore DiO. Relative amount of DiO uptake measured for both micelles. Results represent mean ± SD for n = 5. * = $p \leq 0.05$. Statistical analysis: two-tailed unpaired t-test.

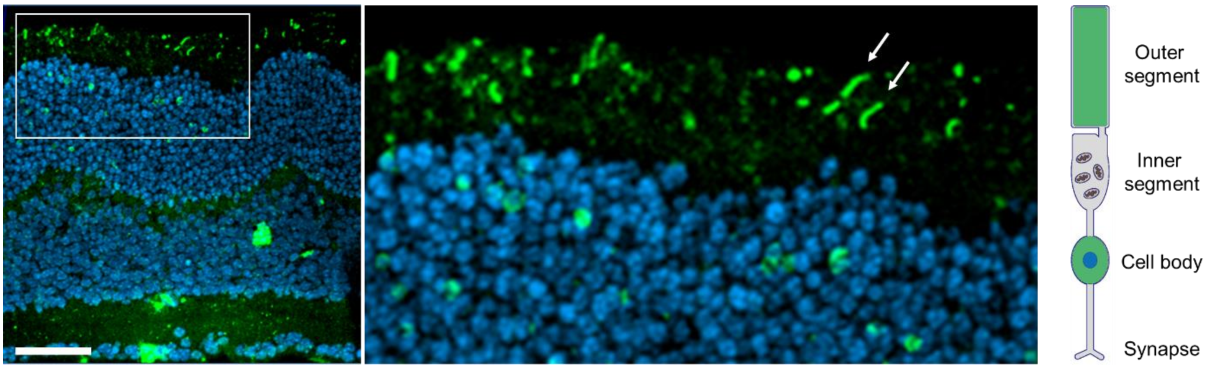


**Figure S2: Intracellular calcein distribution in photoreceptors.** Section from organotypic retinal explant culture derived from wild-type mice to which calcein-loaded pyruvate-liposomes were added. Elongated structures in the outer part of the tissue section most likely indicate intracellular calcein in photoreceptor outer segments (arrows). A qualitative analysis from close-up images revealed high calcein signal in the photoreceptor cell body and outer segments. Scale bar = 50 µm.
